# A survey of mapping algorithms in the long-reads era

**DOI:** 10.1101/2022.05.21.492932

**Authors:** Kristoffer Sahlin, Thomas Baudeau, Bastien Cazaux, Camille Marchet

**Affiliations:** Department of Mathematics, Science for Life Laboratory, Stockholm University, 106 91, Stockholm, Sweden; Univ. Lille, CNRS, Centrale Lille, UMR 9189 CRIStAL, F-59000 Lille, France

**Keywords:** Mapping, alignment, minimizers, syncmers, dynamic programming, seed-and-chain, long reads, third generation sequencing

## Abstract

It has been ten years since the first publication of a method dedicated entirely to mapping third-generation sequencing long-reads. The unprecedented characteristics of this new type of sequencing data created a shift, and methods moved on from the *seed-and-extend* framework previously used for short reads to a *seed-and-chain* framework due to the abundance of seeds in each read. As a result, the main novelties in proposed long-read mapping algorithms are typically based on alternative seed constructs or chaining formulations. Dozens of tools now exist, whose heuristics have considerably evolved with time. The rapid progress of the field, synchronized with the frequent improvements of data, does not make the literature and implementations easy to keep up with. Therefore, in this survey article, we provide an overview of existing mapping methods for long reads with accessible insights into methods. Since mapping is also very driven by the implementations themselves, we join an original visualization tool to understand the parameter settings (http://bcazaux.polytech-lille.net/Minimap2/) for the chaining part.

## 1 Introduction

With the introduction of PacBio long-read sequencing and later Oxford Nanopore Technologies emerged a need for mapping long and noisy sequencing reads. The data proposed new computational challenges of mapping millions of sequences, initially at expected error rates of 10-20%. From the start, authors noticed that the seed-and-extend paradigm used in short-read mapping was not practical for long-reads. First, seed-and-extend would usually rely on a single match before extending, while long-reads required multiple consistent matches along the read to be confidently mapped. Second, the extending part, which relies on alignment algorithms with quadratic time complexity, had to be avoided given the combined length and the frequent insertions and deletions in such data. Early on, the computational problem was compared to whole-genome alignment, with the additional complexity of high error rates. Such observations lead to the novel *seed-and-chain* paradigm for mapping long-reads (see Figure 1). However, the first long-read alignment algorithms using older seeding techniques designed for generic sequence alignment (e.g., BLAST) were not time-competitive in their throughput compared to short-read mappers. Thus, sketching and subsampling techniques imported from comparative genomics started to appear in this domain.

**Figure 1.**
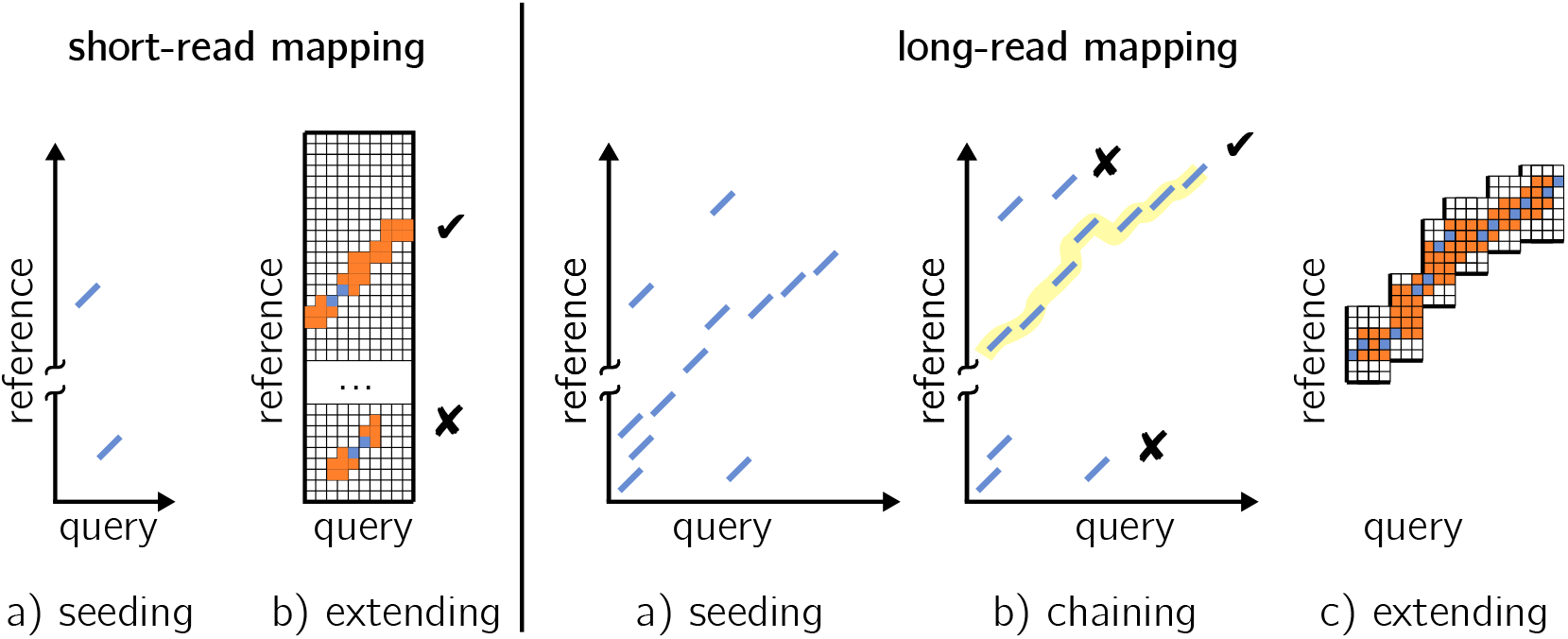
Differences in the main steps between short-read mapping (left) and long-read mapping (right). *Query* denotes the read and *reference* denotes a genome region. Mainly, short-read approaches extend (orange parts) from a single anchor (in blue) on the whole read length while long-read approaches gather multiple anchors, and chain (yellow line) them in for a candidate extending procedure that is done between pairs of anchors.

Recently, specific sub-problems in the mapping domain have been identified and investigated, such as partial and gapped alignment of reads for structural variant discovery, aligning reads in repetitive regions or from non-reference alleles to correct loci, and other applications such as spliced-mapping of RNA reads. These specific problems require and motivate novel algorithmic solutions. In this survey article, we give an overview of the techniques that have been proposed over the last decade for mapping long reads to genomes. After giving definitions and main intuitions, we describe the methodology in two steps. First, seeding, up to the latest advents using novel seeds (syncmers, strobemers). Second, chaining, for which we decipher the currently used score functions. We also made available an original visualization tool to play with the different parameters and to understand their impact on the chain (http://bcazaux.polytech-lille.net/Minimap2).

## 2 Definitions and state-of-the-art of tools

### 2.1 Preliminaries

In this article we restrain ourselves to the problem of mapping a sequence shorter or equal to a genome (a read) to a reference genome. We further assume that the reads come from a genome that is closely related to the reference genome, such as from the same organism or a closely related species.

Let *q* = (*q*_1_, … *q*_*l*_) be the read sequence of size *l* and *t* = (*t*_1_, … *t*_*n*_) the sequence of the reference region of size *n*. Let Σ = {*A, C, G, T* } and Σ_+_ = {*A, C, G, T*, −} be two alphabets, *x* and *y* strings are defined on Σ. Let 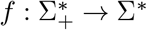 be a transform that maps a string to its subsequence with all “−” characters removed. An alignment is a pair of strings 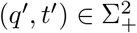 such that:

1. *q*′ and *t*′ have the same size: |*q*′| = |*t*′| = *S*
2. the initial sequences are retrieved through the transform: *f* (*q*′) = *q* and *f* (*t*′) = *t*
3. any pair of characters can be matched at a position *i* of the strings but two dashes: (*q*′[*i*], *t*′[*i*]) ≠ (−, −), for 0 ≤ *i < S*

Many alignments exist for a given pair of strings, in theory, the methods described hereafter aim at finding *good* alignments, i.e. alignments that optimize some distance between the pair of strings. The distance is computed using score functions which give rules on the characters pairing.

With *read mapping*, we mean the procedure to find a read’s location on the reference genome. Typically, long-read mapping is performed by seeding and chaining the seeds into high-scoring regions on the genome. In this study, a *read alignment* implies both that the read has been mapped to a location, and that a pairwise alignment between the read and the genome at the mapped location has been performed. Algorithms exist to compute optimal semi-global pairwise alignments with respect to a score function. However, their complexity in 𝒪 (*n* × *l*), disqualifies them in the context of handling big data such as sequencing data. Therefore, methods of the literature use heuristics to perform read mapping on a reference. They do not guarantee to find the optimal solution.

In our survey, we discuss read mapping to a genome sequence. We will use the terms *query* for a read and *reference* to denote the genome.

### 2.2 Overview of fundamental ideas

To our knowledge, the first mapper explicitly written for long-reads was `BLASR` [12], although short-reads mappers had been adapted for the long-read usage [38, 42, 48]. While solutions specialized for either Nanopore [5] or PacBio [29] characteristics appeared, most modern mappers work for both technologies with adapted parameters. `BLASR` presented itself as a hybrid approach descending from both genome to genome alignment methods (such as `MUMmer` [16]) and short-read mappers. The paper contains seminal ideas used in modern long-read mappers such as the seed-and-chain paradigm.

#### Seeding

Seeding is the first operation in the heuristics used by mapping techniques.

##### Definition 1.

*A* ***seed*** *is a subsequence extracted from the query or the reference*.

The purpose of seeding is to find relatively small matching segments between the query and the reference that serves as markers for reference regions that potentially are similar to the read. The reason seeding is used is that it is typically computationally efficient to find matching seeds that can narrow down regions of interest compared to, *e*.*g*., global alignment of the read to the reference. As we will see in Section 3.1, seeds can be of different nature. Seeding relates to pattern matching, although in sequence bioinformatics, practically all approaches work under the paradigm which indexes the reference and query the index to find matches. The underlying assumption is that once the index is created, it can be used several times to map different query sets. To save space, reference indexes can be in a compressed form. Once matches are found, a second operation aims at finding sets of concordantly ordered seeds between the query and the reference (*chaining*; section 3.3) and to “fill the gaps” between seeds as well as providing the final nucleotide level alignment (*extension*; section 4). Seeding was quickly identified as a critical phase in long-read mapping, which led to novel proposals [50, 43, 72].

#### Sketching and subsampling

An important idea for seeding is *sketching* that was introduced in `MHAP`, a long-read overlap finder implemented in an assembly algorithm [7]. Although long read mappers had already been proven faster than alignment approaches [72], the rationale was to improve the time efficiency of the long-read mapping problem in comparison to the throughput of the second generation sequencing mappers. Sketching consists of compressing the information of a set (here a set of *k*-mers) into a fixed-length vector (a sketch) of representative elements called fingerprints. By comparing two sketches, one can approximate a similarity estimation of the two sets quickly and independently of their initial set sizes. Several approaches exist [9, 55, 14]. MinHash [9] is a sketching technique based on locally sensitive hashing, which produces an unbiased estimator for the Jaccard distance between two sets by selecting a subsample for each set and comparing them in a very efficient way. `MHAP` relied on sketching with this MinHash approach. Thus, `MHAP` overcame a space limitation of `BLASR` which would index the whole reference. The type of matches (exact, fixed-size) induced by `MHAP`’s approach also allowed to perform rapid queries. An important limitation of `MHAP` was that the sampling technique gave no guarantee to uniformly cover the query’s sequence. This led to the development of subsampling techniques which have been adapted to approximate distances between sequences, starting with `minimap` [43]. Seeding is still an active research area of long-read mapping with several recent developments [36, 71, 23, 61]. Sketching and subsampling are discussed in Section 3.1.2.

#### Chaining

A key intuition is that in short-reads mapping, the extending procedure could start after finding a single shared seed between the query and the reference, called anchors (for details on techniques related to the previous sequencing generation, we refer the reader to a methodological survey of short-read mapping [3]).

##### Definition 2.

*An* ***anchor*** *is a matching seed between the query and the reference. It is represented by a pair of coordinates on the query and the reference*.

In the literature, an anchor can also be called “a fragment” or “a match". In long-read mapping, the length of the reads and the short seed length used due to the initial high long-read error rates can lead to a large number of seed matches. It is therefore necessary to reduce the search space by selecting subsequences of ordered anchors (chains).

##### Definition 3.

*Let* 𝒜 = [*a*_0_, *a*_1_, …, *a*_*k*_] *be an list of anchors defined by their coordinates on the reference and the query. A* ***chain*** *is a subsequence of* 𝒜 *of length c* ≤ *k. A colinear chain is a subsequence of* 𝒜 *in which anchors are sorted by such that if i < j, a*_*j*_ *is above and to the right of a*_*i*_ *in the* (*ref erence, query*) *plane*.

Drawing inspiration from genome-wide mapping, `BLASR` introduced a chaining step which aims at selecting high-scoring chains from a set of candidate chains. Chaining allows to reduce the final step of a long-read aligner (the base level extension) to alignment of sub-regions between ordered anchors in chains. Chaining in long-reads has been solved using various dynamic programming procedures [72, 62, 44]. In particular, the continuous work effort put in `minimap2` [43, 44, 45] in both seeding and chaining processes made it a baseline for many other tools’ development. Figure 6 shows the different algorithmic choices over time for seeding and chaining.

While this survey covers the genomic mapping aspects, other important contributions have dealt with adapted procedures in the case of long-read RNA mapping [54, 66, 51, 75], and structural variant identification [69, 49, 25, 74], or other specialized problems [56]. Other related research focused on read-to-read overlap detection [20, 76]^2^, or alignment-free/pseudo-mapping approaches [34, 13]. Finally, here we describe algorithmic solutions working on the nucleotide sequence, but raw signal mappers for Nanopore long-reads is also an active area of research [30, 77, 39].

In the following, we hardly elaborate on complexities for the different algorithms. Some are yet unknown, but in many cases implementations simply use heuristics so that each step’s time is expected to be linear.

## 3 A survey of algorithmic steps

### 3.1 Seeding almost always uses sampled, exact, fixed-length matches

Seeding is the procedure that consists in collecting a set 𝒮 of seeds from the reference, then finding matches between the query’s seeds and 𝒮. In order to find matches efficiently, 𝒮 is stored using an index data-structure. In the following we detail the different types of seeds that can be encountered and

#### 3.1.1 *k*-mers

Substrings of length *k*, or *k*-mers, are perhaps the most commonly used seed in bioinformatics. Such seeds can be extracted from the reference and stored for queries with little computational cost. This makes *k*-mers popular in mapping and alignment applications that require high-performance to scale for millions to billions of reads. A *k*-mer seed can be indexed by using a hash function to produce an integer value (usually as a 32 or 64-bit integer), which is then added to a hash table. This makes indexing of *k*-mers computationally cheap, provided that the hash function and hash table implementations are efficient. Methods to efficiently hash *k*-mers have been proposed [57], which uses the previous *k*-mers hash value to compute the next one using a rolling hash function.

Both a strength and a weakness with *k*-mers are that if a *k*-mer match is found, it is guaranteed to be exact. While it is desirable to produce matches only to identical regions, a downside is that mutations will “destroy” the *k*-mers in the region. By destroy, we mean that *k*-mers different from those of the original sequence are produced, destroying the possibility of exact matches. Typically, a single substitution destroys 2*k* − 1 *k*-mers. This has been studied theoretically in [6] where the authors derived analytical expressions for the mean and variance of regions without matches for a given mutation rate.

#### 3.1.2 *k*-mer subsampling techniques

As any two consecutive k-mers share most of their sequence and are therefore mostly redundant, we could reduce the memory overhead and query time without losing much of the information if not all adjacent or nearby k-mers were stored. In the following, we present different methods that allow picking a subsample of representative *k*-mers as seeds. These approaches have proven their efficiency at reducing drastically the number of objects to index while keeping high sensitivity and specificity for matches.

##### Properties of subsampling techniques

Informally, we define the density of a subsampling technique as the expected number of selected *k*-mers over the total number of *k*-mers [71]. Density depends on the parameters used to select *k*-mers. Aside from the amount of selected *k*-mers, one can be interested in their distribution on the given sequence. The *distance guarantee* (or *r*-window guarantee [71]) ensures that for *r* consecutive positions on a sequence, at least one *k*-mer will be selected by the method. It follows that a method with a distance guarantee has a controlled maximum gap size between two *k*-mers. The choice of representative *k*-mer may or may not be dependent on the surrounding *k*-mers. This property is formalized in the concept of *context dependency* [71]. In context-dependant methods, a *k*-mer’s selection depends on the other *k*-mers nearby. In context-free methods, each *k*-mer’s capacity to be selected is independent. Being context-free implies better conservation of the overall sampled region under mutations. Indeed, context-dependent representatives can tend to be broken over several consecutive windows because of the *k*-mers propagating an error. Finally, other aspects can be considered, such as the deviation of minimizer-based strategies from the initial unbiased Jaccard estimator [6].

##### No distance guarantee between seeds: sketching

Sketching gives typically no guarantee of distance between representatives, which means that a very large gap can appear between two consecutive selected *k*-mers. An early work [7] bases its long-read mapping strategy on MinHash sketching by using a total ordering on the *k*-mers’ hashes, and keeping minimal hashes in the ordering (representing their *k*-mers). Related to read mapping, it was used to perform genome-length sequences alignment-free mapping [34] and to find read-to-read overlaps in long-read assembly [70]. However, fixed-size sketches do not adapt well to different read lengths since the number of fingerprints remains constant for any distinct *k*-mer number. Because of this, two similar regions from sequences of different sizes will not automatically have the same representative, which is a desired property for seeding. Therefore this approach was later replaced by other subsampling strategies in following papers.

##### Distance guaranteed between seeds

On the contrary to sketching, subsampling techniques have been proposed to guarantee that for a certain amount of consecutive *k*-mers, at least one will be selected. The first *k*-mer subsampling technique proposed in the context of long-read mapping was *minimizers*. Minimizers have been introduced in two independent publications [63, 68]. In our framework, minimizers are sampled *k*-mers given three parameters *m, w* and *h. h* is a function which defines an order, which could be the lexicographical order. Given the set *k*-mers starting in a window [*m, m* + *w* − 1] of *w* positions on the sequence, a minimizer is the minimal value for *h* over this set (and therefore the *k*-mer associated with this minimal value) (see left panel in Figure 2). Minimizers are produced by extracting a minimizer in each window *w* ∈ [0, |*S*| − *w* + 1] over a sequence *S*. The techniques used for assigning values to *k*-mers are discussed in section 3.2.1. Minimizers are context-dependent since the content of a window impacts the *k*-mer selection. They have a distance-guarantee property since at least one minimizer is selected by window. Minimizers are agnostic to their relative abundance over a sequence. Different optimizations have been proposed to reduce the density of sampled minimizers in some regions. Weighted minimizers [36] implement a procedure to select *k*-mers of variable rareness. In order for *k*-mers from highly repetitive regions not to be as likely as others to be selected, it first counts *k*-mers, and downweights frequently occurring ones. Then it takes this weight into account for the hashing procedure. Other subsampling techniques include syncmers [18] and minimally overlapping words (MOW) [24]. The first was used in the context of long-read mapping [71] in an alternative implementation of `minimap2` and even more recently in [61]^3^. For their construction, syncmers use *s*-mers of size *k* − *s* + 1 (*s < k*) occurring within *k*-mers (see right panel in Figure 2 for an illustrated difference with the minimizers). The *k*-mer is selected if its smallest *s*-mer meets some criteria. An example criteria is that the *s*-mer appears at position *p* within the *k*-mer (0 ≤ *p < k* − *s* + 1) (these are called *open syncmers*), a more studied category is *closed syncmers* where *p* must be the first or the last *s*-mer position in the *k*-mer. This way of selection uses properties intrisic to each *k*-mer, therefore is context-free. Closed syncmers also have a distance guarantee. By construction, syncmers tend to produce a more even spacing between sampled seeds while still allowing a distance guarantee.

**Figure 2.**
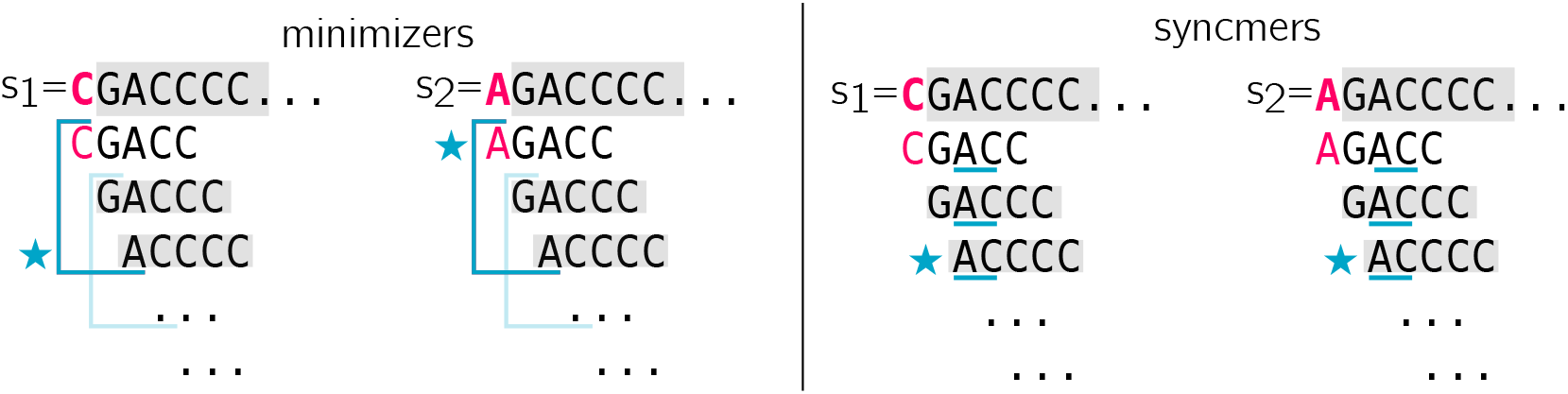
Illusration of minimizer selection, syncmer selection and window dependency. Here two sequences *s*_1_ and *s*_2_ are different from a single mutated base (first base, in pink). When comparing those sequences, one would like to focus on common bases, i.e. bases highlighted in grey. In the left panel, we present a selected minimizer with *k* = 5, *w* = 3. One blue window is presented, a second is suggested in lighter blue. A star shows the position of the selected *k*-mer in the window (we use lexicographic order). The mutated base has an impact on the overall window content, therefore a *k*-mer from the (unmutated) region of interest in *s*_1_ is no longer selected in *s*_2_. On the contrary, in the right panel, we show that syncmers can be more robust in this situation. We choose *k* = 5, *s* = 2 and present closed syncmers. We underline the smallest *s−*mer in each *k*-mer in blue and a star shows the selected *k*-mers. We see that in this example, the mutated base has no impact on the syncmer selection, and the same syncmer is selected in the region of interest for *s*_1_ and *s*_2_.

#### 3.1.3 Fuzzy seeds handling substitutions

Due to read errors and SNPs between the reference and sequenced organism, it is in many scenarios desired that a seed match between the query and the reference even if the seed contains a substitution. Put differently, we would want similar *k*-mers to hash to identical hash values. A hash function that would produce identical hash values for similar but not necessarily identical inputs is usually referred to as a locality-sensitive hash function. We will refer to seeds produced under such methods as fuzzy or inexact seeds.

Several methods to produce inexact seeds have been described. Perhaps the most common one is spaced-seeds. Within a spaced-seed, some positions are required to match (called fixed positions) while the remaining positions can be ignored (called wildcards or don’t care positions). Within a *k*-mer, fixed positions can be selected to be wildcards by applying particular masks on the *k*-mer’s bases [33]. A problem with spaced-seeds is to find a fixed-position profile to minimize the overlap of the fixed positions in the seeds [32]. Although the computation of *good* spaced-seeds has been optimized [33], constructing these spaced-seeds profiles requires extra computational work compared to *k*-mers and is therefore slower to compute, and in practice, multiple different seeds are used [47] to increase sensitivity, which requires storing multiple hash tables. Another limitation with fuzzy seeds for substitutions is that seeds will, just as for *k*-mers, not match over indels.

While fuzzy-seeds handling substitutions have been used e.g. in metagenome short-read classification [10] and permutation-based seeds were implemented for short-read mapping [41], few of long-read mapping algorithms implement them. As indels are a frequent source of variability on long-reads, the computations to construct these seeds may not be worth the trade-off in increased sensitivity. An exception to this is a recent seeding mechanism [23], where the authors use a variant of SimHash [14](an alternative locality sensitive hashing to MinHash) to construct fuzzy seeds over subsampled *k*-mers using the minimizer technique [63]. The authors showed read alignment can be improved both in terms of speed and accuracy by integrating their seeds into `minimap2` [44].

#### 3.1.4 Fuzzy seeds handling indels

A common source of errors and biological variation is short insertions and deletions. Neither the exact seeds nor the fuzzy seeds discussed so far are designed to match over such variability. Traditionally, matching over indels has typically been solved not by a single query of a fuzzy seed, but instead involved queries of a few short *k*-mers at a close occurring distance which are then inferred as a matching region. While several queries in a nearby region usually provide gold standard sequence similarity queries [4, 37], it comes at a significant computational cost. Along the same vein [72] proposed to index one so-called spaced *k*-mer as a seed in each position of the reference and would, query three different seeds for each position in the query (representing a mismatch, a deletion of length one, and a mismatch and a one nucleotide insertion). This design was motivated by overcoming the frequent substitutions and short indels present in third-generation sequencing techniques, but would only handle indels of one nucleotide (we provide details on this scheme in Supplementary Figures S1 and S2). Earlier, there have been works to handle higher error rates with so-called covering template families [27] that can guarantee a match up to any error rate *e*. Naturally, with higher *e*, more seeds need to be indexed and queried and it becomes computationally prohibitive to use such seeding.

To remove the overhead of post-processing of nearby seeds [4, 37] or multiple queries [72] per indexed reference seed, one can instead link the *k*-mers up into a seed before storing it in the index. Such indexing has been favorable in the long-reads era where indels are frequent. One proposed method is to join two nearby minimizers into a seed. Joining nearby minimizers is usually a relatively cheap computation as the minimizers constitute a subset of the positions on the reference. Such a seeding technique has been used for long-read overlap detection for both genome assembly [15] and error correction [67]. While such indexing is relatively fast and matches regions over indels, the joining of nearby minimizers implies that if some minimizer(s) are destroyed due to mutations in a region, all of the seeds in that region will be destroyed. Put another way, nearby seeds share the same information (in the form of a shared minimizer). Therefore, alternative approaches such as strobemers [64] (see right panel in Figure 3) have been described, where the goal has been to reduce the information between closeby seeds by linking *k*-mers at seemingly random positions within a window. Such pseudorandom linking implies that if one seed is destroyed due to a mutation, a nearby seed may still match. Strobemers have shown effective at finding matches between long-reads and for long-read mapping [64], and have been used in short-read alignment programs [65] but they come at an increased computational cost to joining neighboring minimizers.

**Figure 3.**
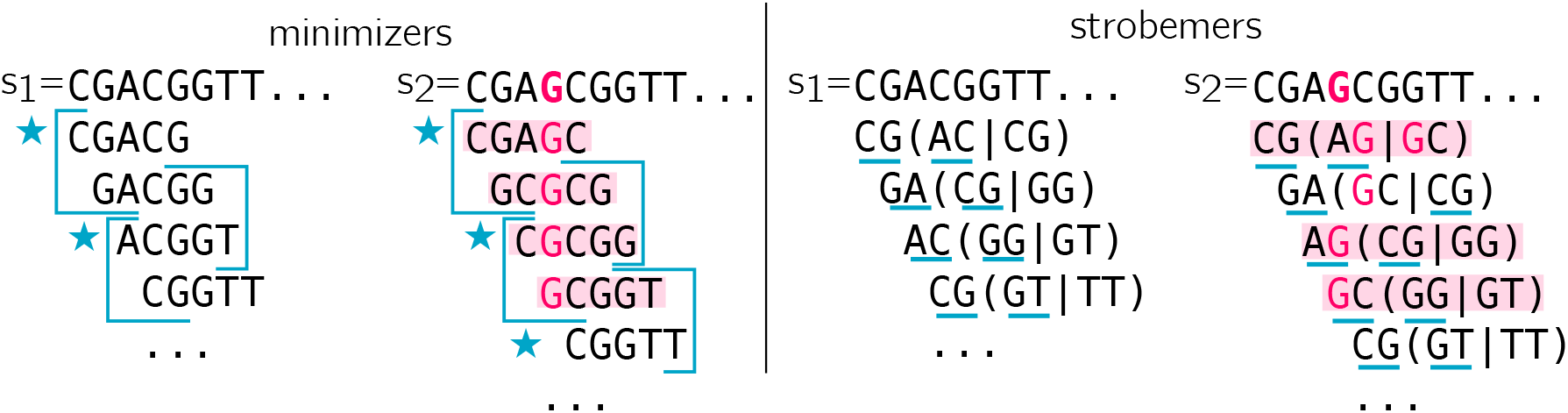
Illustration of strobemers’ capacity to handle indels. As in Figure 2, two sequences are presented. This time, *s*_2_ has an insertion (pink G). On the left panel, minimizers are selected using *w* = 2, *k* = 5. Blue stars point selected minimizers in each blue window. One can see that the only safe region to generate minimizer is the CGGTT sequence after the insertion, that is shared and of length *≥ k*. Put differently, *k*-mers in red have no chance to be in common between the two sequences. However, in this example, the scheme fails to select a common minimizer in the safe region. Strobemer selection is presented in the right panel, using *k* = 2, *s* = 2, *w* = 2. At each position, the first *k*-mer is selected to be the start site of the strobemer. Then, in the non-overlapping window (of size *w*) downstream to the first *k*-mer, a second *k*-mer is selected according to one of the selection techniques presented in [64] (we illustrate selecting the lexicographical minimizer). We underline the bases that are kept for each strobemer. For instance in *s*_1_, the first *k*-mer is CG at positions 0 and 1, then the next window starts at position 2. Two *k*-mers are computed from this window, AC and CG, and AC is the minimizer. Therefore, the strobemer is (CG,AC). Again, strobemers with no chance to be shared between *s*_1_ and *s*_2_ are colored in red. For strobemers, it is the case when at least one part contains the mutated base. We note that not only the CGGTT region has a common strobemer (CG,GT) in both sequences, but also that the scheme allowed to “jump over” the mutated G and could select another common strobemer (GA,CG) in a more difficult region. The strobemers in this example consists of two *k*-mers (*s* = 2) but they can be constructed for other *s >* 2.

Another way to alleviate the issue that mutations will destroy consecutive seeds in the neighboring minimizers technique is to apply the SimHash technique on strobemers instead of *k*-mers [23]. Such seeds were used for long-read overlap detection [23] and the authors show that for the highest quality long-reads (PacBio HiFi), such seeds can speed up long-read overlap detection by an order of magnitude or more while retaining the same downstream level assembly accuracy.

#### 3.1.5 Dynamic seeds

Previously discussed seeds share the characteristic that they can all be produced and inserted in a hash table, and consequently, only require a single lookup. This is typically fast and, hence, popular to use in long-read alignment algorithms. The downside is that if a seed is different in a region between the reference and the query (e.g., due to an error), there is no way to alternate the seeds in this region at alignment time. There are however other types of constructs, that we here refer to as dynamic seeds, that can be computed on the fly at the mapping step, and then used as seeds downstream in the read alignment algorithm.

##### Maximal Exact Matches

Maximal exact matches (MEMs) [16] are matches between a query and reference sequence that cannot be extended in any direction on the query or reference without destroying the match. These are typically produced by first identifying a *k*-mer match, and then an extension process is applied. MEMs are guaranteed to be an exact match between the query and the reference and are bounded below by length *k* but do not have an upper threshold for seed size. As there can typically exist many MEMs, a subset of MEMs that has a unique location on both the query and reference is sometimes considered. MEMs or similar approaches have been used in one of the earlier long-read alignment programs (e.g., `BWA-MEM`) [42, 12] and for long-read splice alignment [66], but these seeds are more computationally expensive to compute and are typically slower than single-query seed-based algorithms.

##### Anchors from minimal confidently alignable substrings (MCASs)

If a query was sampled from a repetitive region in the reference, one may likely find several clusters of anchors across the reference. Further dynamic programming operations to decipher the true origin region of the query are typically costly or even unfeasible if too many copies have to be considered. Even in the case a query is located on the reference, it might be attributed to the wrong copy because of the sequencing errors. A recent contribution [35] proposed a solution for handling seeding in repetitive regions. The procedure finds smallest subsequences that *uniquely* match (MCASs) between the query and the reference. There can be as many as the query length in theory. In practice, the more the repeats are divergent, the shortest the MCASs since a base pertaining to a single copy is more likely to be met. MCASs are computed using an alignment procedure, which means that *uniquely* matched must be understood as a relative property. For each position on a query, the best and second-best alignment scores are compared, and a substring is considered uniquely matched if the difference between the scores is above a threshold. It is interesting to bound the maximal size of MCASs, both for performance purposes and because they may become less specific as to their size increase. Fixed-size, exact match anchors (minimizers) are then extracted from MCAS regions.

### 3.2 Implementation of the seeding step

#### 3.2.1 Seeds transformations before indexing

Originally, minimizers use a lexicographical ordering. However, in our four base alphabet, this can tend to select sequences starting with long alphabetically smaller runs such as “AAA… “. Random hash functions assigning each *k*-mer a value between 0 and a maximum integer are preferred [68].

Oxford Nanopore reads are known for accumulating errors in homopolymers, typically adding/removing a base in a stretch of a single nucleotide. Sequences can be homopolymer-compressed before finding *k*-mers. Homopolymers longer than a size *s* are reduced to a single base, then *k*-mers are computed over the compressed sequence. For instance, for *s* = 3, *k* = 4, an original sequence ATTTTGAAAACC is compressed to ATGACC, and the final *k*-mers are ATGA, TGAC, GACC. This procedure allows finding more anchors while indexing fewer *k*-mers/minimizers. Homopolymer compression is ubiquitous in long-read mappers.

In regions of low complexity (e.g. ATATATA, CCCCC) the standard minimizer procedure keeps all minimal *k*-mers in windows. It is then possible for two *k*-mers to get the minimal value and to be selected, which tends to over-sample repetitive *k*-mers. A *robust winnowing* procedure is proposed in [36], which avoids the over-sampling effect by selecting fewer copies of a *k*-mer, but increases the context dependency phenomenon.

#### 3.2.2 Hash tables prevail for seed indexing

Indexing of fixed size is usually done using hash tables (although FM-indexes for *k*-mers exist [8]). In the context of subsampling, invertible hash functions have been a key asset for using minimizers as *k*-mers representatives. In other words, a hash value is associated with one and only one *k*-mer, and the *k*-mer sequence can be retrieved from the hash value (using reciprocal operations). This choice allows a very fast *k*-mer/minimizer correspondence but is costly as it implies that the fingerprints of the hash table are not compressed (which is mitigated by the subsampling). Minimizers are then used to populate a hash table, which associates them to their position(s) in the reference and their strand information (usually hashed seeds are canonical *k*-mers: the smallest lexicographic sequence between the original *k*-mer and its reverse complement).

Variable-length seeds are indexed in full-text data structures (suffix arrays, FM-index [22]), which allow to find and count arbitrarily long queries in the reference. They have been used in the first versions of long-read mappers. Variable-length seeds type can be longer to query in the structure, while hashed matches are queried in constant time. Since minimizers represent fixed-length *k*-mers, hash table solutions mainly prevail.

#### 3.2.3 Seeds selection at the query

In [44], it is proposed to select all minimizers from the reference during the indexing phase (although the latest versions include the weighted *k*-mers and robust winnowing heuristics), and to soft mask some representative *k*-mers at the query. The procedure simply avoids *k*-mers seen too many times according to a fixed cutoff. The authors noticed that in cases where a query is sampled from a repetitive region, such a procedure prevents it to be seeded. Therefore, an update was proposed [45], which detects if low occurrence *k*-mers are too far away in a query, and in this case, allows sampling minimizers in the repetitive region in between (and keeps some of the lowest possible occurrences among these minimizers). Techniques that use longer fuzzy seeds (e.g., strobemers) [23] reduce the number of masked regions, although it comes at the cost of sensitivity. Another approach [62] computes a new set of minimizers on the targeted reference region in order to obtain finer candidate chains, in particular in repeated or low complexity regions.

### 3.3 Chaining is dominated by dynamic programming with concave gap score functions

#### 3.3.1 A dynamic programming problem

Once the reference’s seeds are indexed, a set of seeds is extracted from the query and looked up in the index to find anchors. Anchors’ positions on the query and reference are stored, as well as the forward/reverse information. Instead of directly extending the alignment between anchors, as it is done in short-read mapping, a step of chaining is added and meant to accelerate further extensions. Chaining acts as a filter and a guide for smaller extensions that need to be realized only between selected anchor pairs. Without it, too many extension procedures, most of which would be dead-ends, would have to be started.

In an ideal case, there is a unique way of ordering anchors by ascending Cartesian positions in the (*reference, query*) space, which passes by all the anchors. In practice, some anchors are spurious, others correspond to repeated regions and yield different possible chains. Moreover, over parameters have to be taken into account. Thus, methods optimize different aspects (also illustrated in Figure 4):

**Figure 4.**
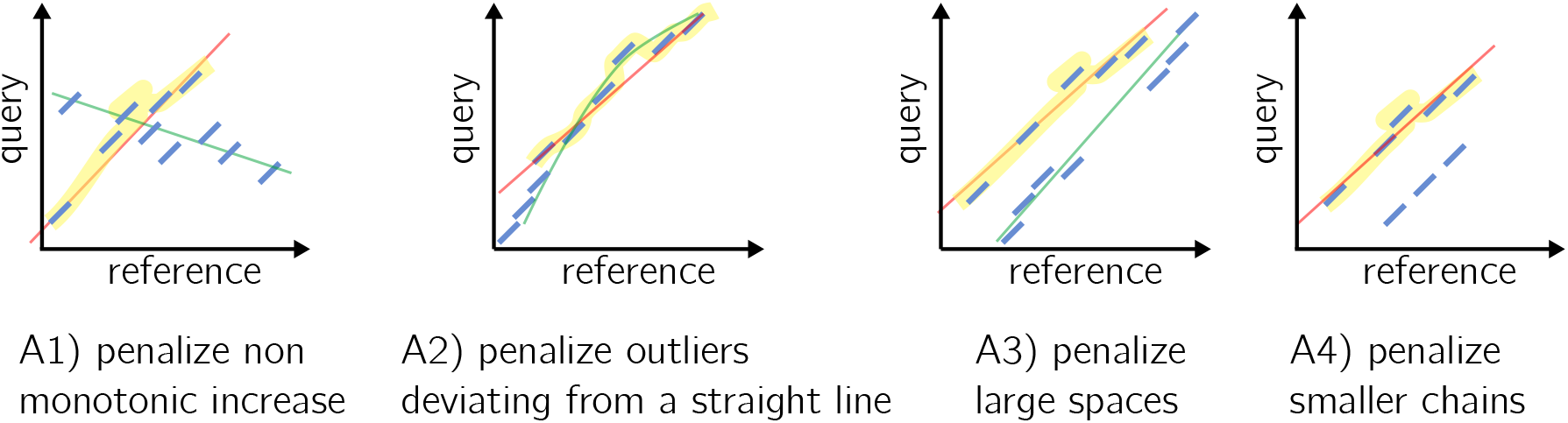
An illustration of the different constraint taken into account in the gap score functions. The reference axis shows a genome region of interest where anchors were found, not the whole reference. A1–A4 correspond to items in the text in section 3.3.1. Anchors are showed in blue. The selected chain with respect to the described constraint is highlighted in yellow and a line approximately passing by its anchors is showed in red. The line passing by the longest chain is showed in green.

A1) Do not allow anchors which are not ascending either by the anchors’ start or end coordinates in both the query and reference (see first case in Figure 4).

A2) Avoid discrepancies in diagonals between anchors (second case in Figure 4).

A3) Do not allow large spaces between consecutive anchors of the chain (see third case in Figure 4).

A4) Favor the longest possible anchor chain (fourth case in Figure 4).

A5) If inexact matches in seeds are possible, find a series of anchors ensuring a minimal Levenshtein distance between the query and the reference.

The problem of finding an optimal chain using non-overlapping anchors has been called the *local chaining problem* [1], although in this application anchors can overlap. The score *f* (*i* + 1) represents the cost of appending an anchor *a*_*i*+1_ to *a*_*i*_ to the chain. This score is often called the *gap score* in the literature, though it includes other constraints, as described above. The chaining problem for long reads seeks to find an optimal colinear chain with a positive gap score.

Mainly, methods use either a two-step approach: 1-find rough clusters of seeds as putative chains, followed by 2-find the best scored chain among the selected clusters; or work in a single pass and apply a custom dynamic programming solution to find the best anchor chain. We can start by noting that one of the first mappers dedicated to long-reads solved a global chaining problem to determine a chain of maximum score, by fixing starting and ending points (anchors) such that their interval is roughly the size of the query [12]. Such an approach would easily discard long gaps and spaces in alignments.

#### 3.3.2 Chaining in two steps

##### Clusters of seeds are found through single-linkage in 2D space

The two-step approaches rely on a first clustering step. Although it tends to be replaced by single-step chaining (see Section 3.3.3), in the following we describe the fundamental ideas of the clustering. Methods first find rough clusters of anchors by considering a discrete (*reference, query*) position space. In this space, an anchor realizing a perfect match is a line of the size of the seed. This line should have a 45-degree angle, which also corresponds to the main diagonal of a (*ref erence, query*) alignment matrix. The same idea stands for a set of anchors. However, because of insertions and deletions, each small line materializing an anchor may not be on the exact same diagonal, thus realizing approximate lines in the (*reference, query*) space. A method from image processing has been proposed to find approximate lines in this space: the *Hough transform* [17], which makes it possible to detect imperfect straight lines in 2D space. Contrary to linear regression which would output the best line explained by the anchor distribution, here an arbitrary number of straight lines can be output and considered (see Supplementary Figure S3 for an illustration). Hough transform or other similar anchor grouping algorithms ([62] proposes to delineate fine-grained clusters in order to increase the chaining specificity in repeated regions) all can be assimilated to single-linkage clustering in 2D space, which finds groups of anchors placed roughly on the same diagonal.

##### Anchor chaining using longest subsequences of anchors

The previous clustering techniques aim at finding lines in groups of anchors that can be approximately colinear. To determine truly colinear chains, a subset of anchors can be ordered by finding a longest increasing subsequence (LIS) of anchors. Let each anchor be mapped to 1 … *n* integers. The LIS problem consists in finding a longest increasing subsequence from a permutation *P* of the set {1, 2, … *c*}, which can be solved in O(*c* × *log*(*c*)).

In the case of exact fuzzy seeds, inexact matches are to be dealt with on top of the initial increasing chain problem. Indeed, one wants to obtain the closest base-wise anchor chain. In this case, the problem is converted to LCSk (longest common subsequence in at least *k*-length substrings). Note that there is a correspondence between LIS and LCS. The LIS of *P* is the LCS between *P* and the sequence (1, 2, … *c*). In both cases, neither the longest nor the increasing requirements are sufficient to find correct anchor chains: they lack definitions for other constraints, such as distance between anchors or the possibility to allow large gaps. They are complemented with heuristics or replaced by more recent approaches in Section 3.3.3. In addition, several methods use graphs built over anchors as backbones to the chaining and alignment steps [74, 50, 72] (one approach is described in the Appendix). Because they would fail to take into account distances between anchors, these methods have been replaced by dynamic programming approaches relying on gap score functions.

#### 3.3.3 Chaining in a single step: gap score functions

The main drawback of the approaches previously described in 3.3.2 is that though large spaces between two anchors of a pair must be avoided, some spaces correspond to gaps in the alignment and can be kept. In order to deal concurrently with these two problems, most recent methods drop the two-step clustering and LIS to directly apply a custom dynamic programming solution. It is globally the same spirit as LIS, but integrates a more fine-grained gap penalty solution. It defines a cost function that grants a maximum penalty for non-monotonic increasing seed chains.

##### Concave gap functions

The cost function is designed to handle the gaps induced by frequent indels in the data. Intuitively, it is likely that indels happen in clusters of *n* positions rather than at *n* independent positions in the chain because some regions on the query might be particularly spurious, or because of local repeats on the reference. Therefore, the same cost is not attributed to opening a gap and extending a gap, thus a linear gap function does not fit. The choice of gap functions which are concave (verifying the *Quadrangle Inequality*) improves the time complexity by using the *wider is worse* strategy [26, 21]. In practice, these concave gap functions are affine, a set of affine functions, or a combination of affine and log functions, as proposed in [44]. We chose to present `minimap2`’s [44] gap functions in Figure 5 as they are adopted without modifications in most current papers (with the recent exception of [62]). Chains are built by aggregating close anchors of smaller coordinates to the current anchor by penalizing the shifts compared to the main diagonal. In Figure 5, Panel 5a presents how the set of possible anchors to prolong the chain is selected. Panel 5b illustrates the dynamic function’s parameters. The complete description of the functions is available in the Appendix. Here we recall the formula for regular gap sizes, for anchors *i* and *j*:

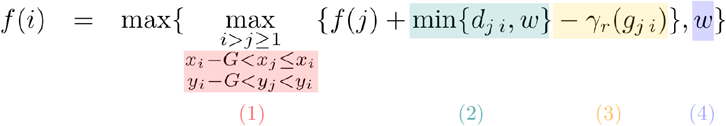

**Figure 5.**
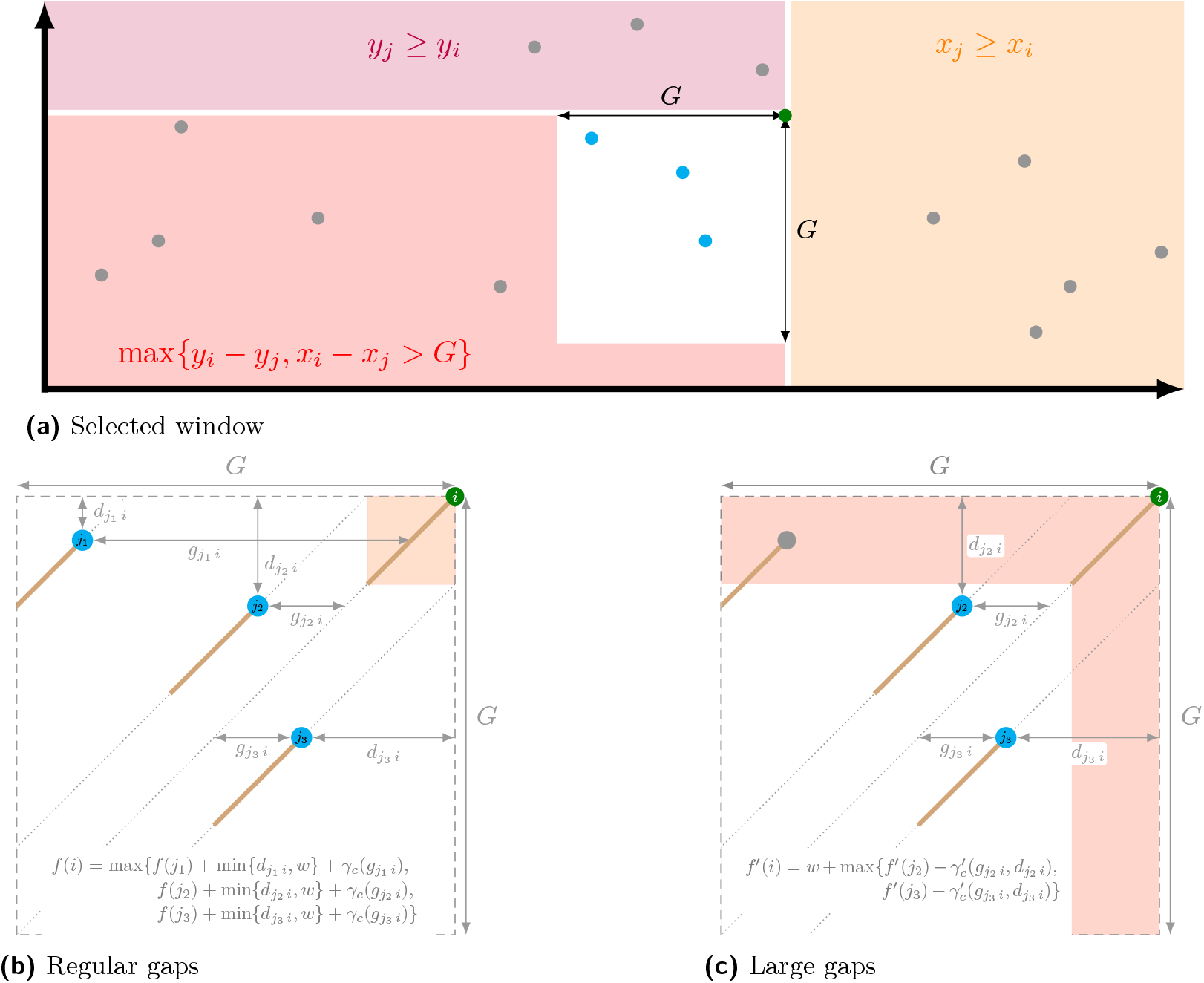
Outline of `minimap2`’s chaining. Figure a shows for an anchor (in green) the selected region (in white, *G* is the gap threshold) to find available anchors to continue chaining (in blue). Figures b and c give respectively the dynamic programming functions for regular and large gaps size. Anchors are shown as segments ending with green or blue dots with the same color code as in Figure a. Besides, for the large gap size (Figure c), to improve the complexity, the anchors do not overlap (available anchors are not in the red zone). *d*_*ji*_ represents the smallest “distance” between the two anchors (but is not really a distance by definition), *w* is the minimizer window size, *g*_*ji*_ is the gap length, and the *γ* functions are the concave gap functions.

**Figure 6.**
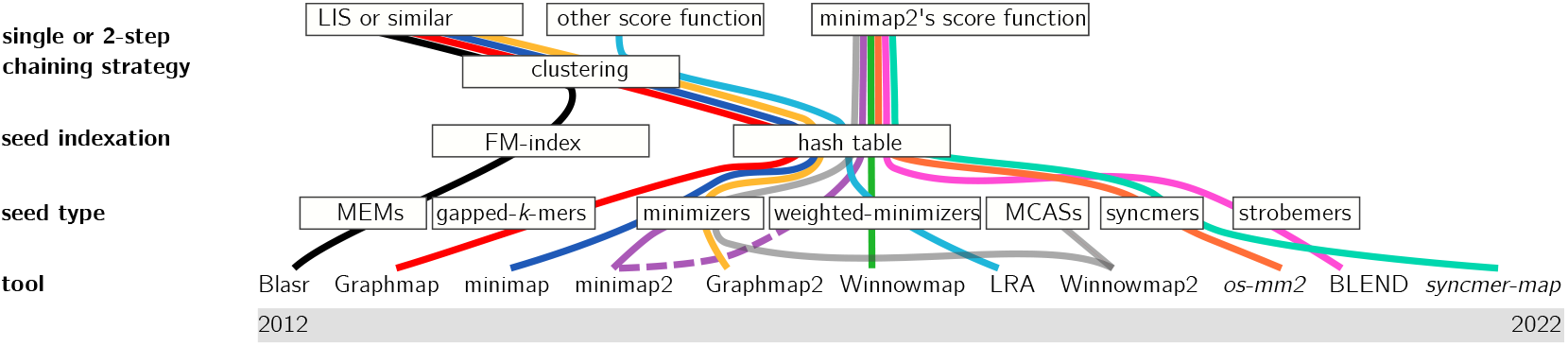
Long read mapping tools over time. Tools and techniques are presented from oldest to most recent, from left to right. The figure presents implementation names at the bottom, then goes up to the different steps: seeding, with seed selection strategies and indexation, then chaining strategies. Alignment is not represented as it is not very different from one tool to another. The dotted line for `minimap2` means its implementation evolved. The italic names denote for a proof-of-concept rather than an tool.

With *f* (*i*), *f* (*j*) scores for anchors *i* and *j*, and:

(1) This property means that at least on axis X or Y, the distance between the two anchors must be *> G* the maximum authorized gap size, and that *i* is above and to the right of *j* (see the three colored zones in panel (a) of Figure 5).

(2) This minimum penalizes overlaps between anchors. Indeed, if *j* starts before a distance of *w* on the diagonal (with *w* the minimizer window size and window guarantee of minimizers) then *d*_*ji*_ (the smallest coordinate difference on either X or Y axis between *i* and *j*) will be lower than *w* and therefore selected. For any other case, *w* which is a larger value is selected and increases more the score.

(3) This is the concave gap score penalty (see the Appendix for a complete formula). It is computed on the gap length *g*_*ji*_ = |(*y*_*i*_ − *y*_*j*_) − (*x*_*i*_ − *x*_*j*_)|. It penalizes distant anchors in the Manhattan definition, i.e. anchors far from *i* either on the main diagonal or because they are on side diagonals (see for instance anchor *j*_1_ in panel (b) of Figure 5).

(4) This term helps with the initialization.

In order to test *w* and *γ*_*r*_’s impact on the chain selection, we implemented a visualisation tool: http://bcazaux.polytech-lille.net/Minimap2/. We generate a scenario of shared anchors between two sequences and allow to set the different parameters’ values. We show the selected chain according to the settings. Heuristics are applied to rapidly drop a dynamic programming procedure in regions that are unlikely to align and to avoid 𝒪 (*c*^2^) worst cases. Based on empirical results, these heuristics mostly check if seeds are not separated by too large regions and drop the chaining procedure if the score becomes too low.

##### Solutions for large gaps

Noticing that [44]’s original approach would be failing in large gaps, one contribution [62] proposed techniques to perform dynamic programming with a family of concave functions by relying on a previous work [21] (built on a prior clustering step as described in 3.3.2). Recently, [44] integrated a solution designed for mapping long structural variants in pangenomic graphs [46]. Its recent versions entail a cost function for regular gaps, and a long gap patching procedure. Then it chooses the cheapest solution to move on to the alignment step. The gap patching procedure uses a linear gap cost so that it has a higher long-gap opening cost in comparison to the regular procedure but at a cheaper extending cost. The chaining with a linear function is solved with a range minimum query (RMQ) algorithm using a balanced binary search tree [1, 60]. It allows to solve the linear chaining in O(*c* × *log*(*c*)). Although this time complexity can be improved in O(*c*) by using range maximum queue [11], the implemented algorithm is more costly than the solution for regular gaps, which is preferred if possible. Panel 5c in Figure 5 illustrates the dynamic function for large gaps.

#### 3.3.4 Mapping quality scores have been adapted for ranking chains

The described methods may deliver a set of chains that satisfies the chaining score threshold. To choose among the candidates and decide the final location, chains can then be categorized into primary/secondary chains. Chains with a sufficient score are ranked from highest to lowest score. Primary chains are those with the highest scores which do not overlap with another ranked chain for the most of their length. Secondary chains are others. Mapping quality, which is a measure that had been introduced to assess short-reads mapping, is redefined for long-reads with slight variations according to articles. It reports, for chains, whether the primary is very far in terms of score from the best secondary, and if it is long enough.

## 4 Extension step and final alignment computation

### Extenstion step

In order to allow gaps, the methods rely on local alignment between pairs of successive anchors using classical algorithms [28, 58] derived from Needleman and Wunsch [59]. They are based on alignment matrices, which aggregate the base-wise alignment scores from the two prefixes (top left of the matrix) to the two suffixes (bottom right).

To compute the scores and report them in a matrix, affine cost functions allow to allocate different penalties for opening and extending gaps and therefore can favor short or long gaps. More precisely, such algorithms use pairs of affine gap score functions and choose the cheapest cost between the scoring for short gaps (i.e. less costly to open, costly to extend), and the scoring for long gaps (i.e., more costly to open, cheap to extend). Allowing long gaps has a drastic negative impact on the alignment efficiency because more cells in the alignment matrix have to be considered.

#### Heuristics for speed-up and quality enhancement

Therefore, alignment is commonly accelerated through vectorization, using single instruction multiple data (SIMD) sets of instructions, which increase the computational throughput by passing simultaneously several matrix cells for the processors to evaluate. Second, practical alignment implementation relies on banded alignment, which, simply put, bounds the alignment matrix in a band of size *ℓ* around the top-left – bottom-right diagonal. Inspired from `BLAST`’s X-drop [4], [44] implements a Z-drop procedure. X-drop quits extending the alignment if the maximum score reached at some point when aligning the prefix drops by more than X. Z-drop adds the possibility not to drop the extension during large gaps.

Due to sequencing errors, some spurious anchors main remain in a chain, which can lead to a suboptimal alignment. At the alignment step, [44] chooses to remove anchors that produce an insertion and a deletion at the same time (>10bp) or that lead to a long gap at the extremity of a chain. Another solution [12] involves to re-compute a chain with novel anchors computed on a window that comprises the alignment.

## 5 Future directions

On top of mentioned novel seeding techniques bringing new properties concerning theircoverage of the seeded sequence and robustness to errors and mutations (syncmers, strobe-mers [71, 61, 23]), we can expect to see advances in the chaining and extending parts in the coming months.

Indeed, the usage of *diagonal-transition algorithms* which was initially define for edit distance [73, 40, 31] has been reactivated recently for the gap-affine model with the wavefront alignment algorithm (WFA, including [53, 52, 19]). More precisely, instead of using dynamic programming on the adjacent cells, WFA transposes the optimization problem on the diagonals and the score. In particular, WFA has the potential to make computation faster for similar sequences and large gaps (by setting the score accordingly and adapting the scoring). A current result shows that we can exploit the massive parallel capabilities of modern GPU devices to accelerate this wavefront alignment algorithm [2]. Currently, different implementations exist that have been tested on long reads [53]^4^, although no dedicated long-read mapper integrates them yet.

## Acknowledgements

The authors would like to thank Mikaël Salson and Laurent Noé for proofreading the manuscript and suggesting revisions.

## Appendix

### Graphmap’s indexing strategy for fuzzy seeds

Graphmap builds two hash indexes from two types of shapes, called `6-1-6` and `4-1-4-1-4`. As shown in Supplementary Figure S1, for each position is seeded (no subsampling). A seeded key corresponds to the subsequence at a given position of the reference when applying the shape mask: each *don’t care* (*) base is skipped.

Then, for each read in the query, several lookup keys (for mismatch, deletion and insertion) are built from a shape (Supplementary Figure S2). To that extent, the lookup key treats the *don’t care* base in three different ways. The mis(match) shape has the same behaviour as the indexed key, i.e., the *don’t care* base are skipped. The insertion shape skips two bases: the initial *don’t care* base and the base next to it. Finally the deletion shape will simply build the key and keep all the base including the *don’t care* base. In total, for a number *d* of *don’t care* base, 3*d* different keys are built per shape.

**Figure S1.**
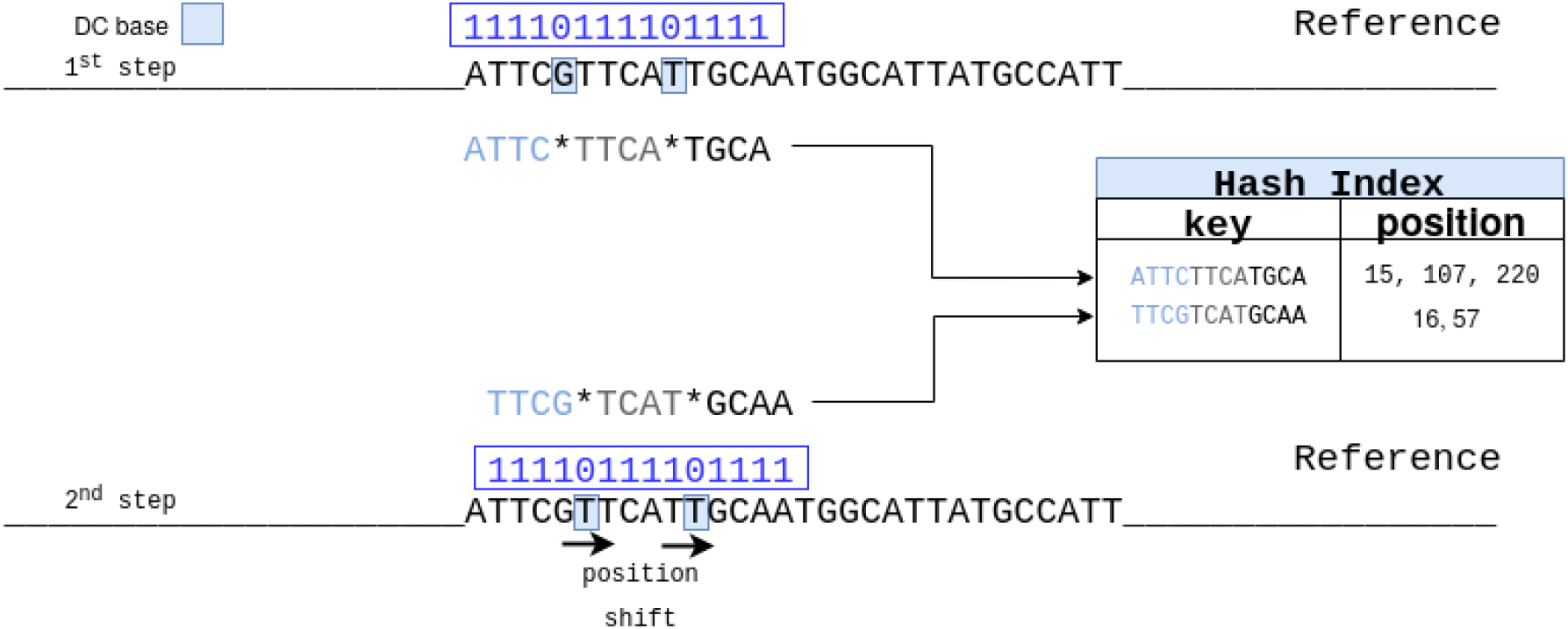
Indexing scheme for fuzzy seeds allowing indels and substitutions in Graphmap. In the figure the shape 4.1.4.1.4 is represented. The zeros represent the don’t care positions of the shape. The shape is then applied for each position of the genome. The substring built from the shape is used as key inside a hash index. Each key will correspond to one or more positions on the reference.

### Graphmap’s backbone graph for LCSk

Because of the possible spurious matches that occur because of the ambiguous bases, `Graphmap`’s fuzzy seeds require more treatments to find proper chains. A first step after seeding finds groups of anchors representing longer shared subsequences between the query and the reference, on which is applied LCSk. Anchors are placed in a vertex-centric positional graph of *k*-mers, in which *k*-mers in both sequences appear, and share an arc if they are directly consecutive (or consecutive up to a distance parameter)5. Most weighted paths of anchors (i.e. supported by the query and the reference) are found in this graph and output as shared subsequences. After the LCSk pass, a L1 linear regression step is applied to fit a straight line with a 45 degree slope and remove outliers, especially in the beginning and end of the chain (see case 4 in Figure 4 in the main text). Note that other methods use graphs built over anchors as backbones to the chaining and alignment steps without fuzzy seeds [74, 50].

**Figure S2.**
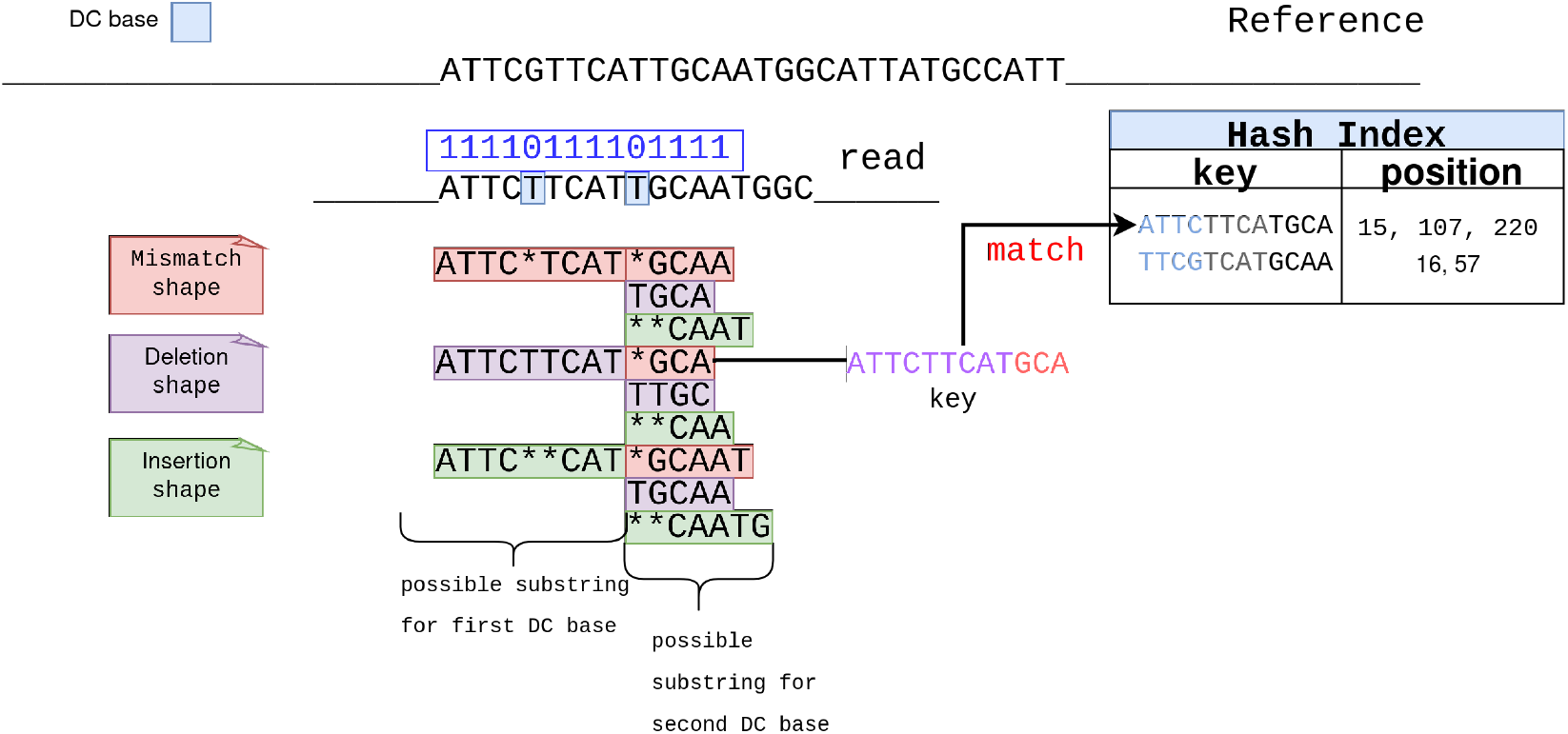
Query in Graphmap, different possible sequences can be matched using a single key. As we can see, there are three types of look-up shapes, and each of them is used to reconstruct a different substring. Each type corresponds to three phenomena that can occur with errors in sequencing, namely substitution, substitution + 1 insertion, and substitution + 1 deletion. Here, two don’t care bases are present and nine substrings can be obtained. In this example the substring obtained from the substitution + insertion shape and the mismatch leads to a match with the reference.

### Minimap2’s complete formula for regular and large gaps size

For regular gaps size:

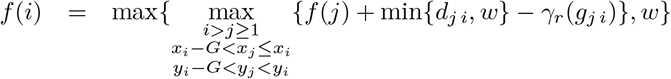

The property *x*_*i*_ − *G* ≤ *x*_*j*_ ≤ *x*_*i*_ and *y*_*i*_ − *G* ≤ *y*_*j*_ *< y*_*i*_ is equivalent to *y*_*j*_ *< y*_*i*_, *x*_*j*_ ≤ *x*_*I*_ and *e*_*j i*_ *< G*.

For large gaps size:

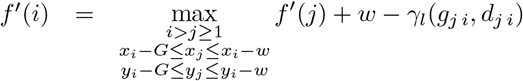

where

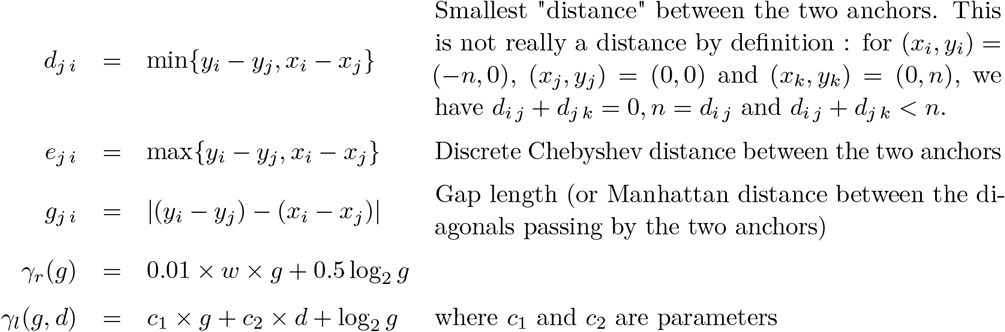

#### Hough transform principle

Applying the Hough transform means going from the *S*1 = (*query, ref erence*) space to the Hough *S*_2_ space of coordinates. If a line (*y* = *ax* + *b*) exists in *S*_1_, it is a point of coordinates (*a, b*) in *S*_2_ (practically, polar coordinates are used for technical reasons). All possible lines intersecting a point in *S*_1_ can be translated in *S*_2_ as a sine wave. Multiple anchors give multiple points in *S*_2_, and the intersection of possible sinusoids intersecting the different points in *S*_2_ correspond to a line roughly intersecting the initial anchors in *S*_1_. The Hough space is rasterized, and by counting and weighing the possible solutions in *S*_2_, lines can be deduced in *S*_1_. Contrary to linear regression which would output the best line explained by the seed distribution in *S*_1_, here an arbitrary number of straight lines can be output and considered (see Supplementary Figure S3 for an overview of the steps).

**Figure S3.**
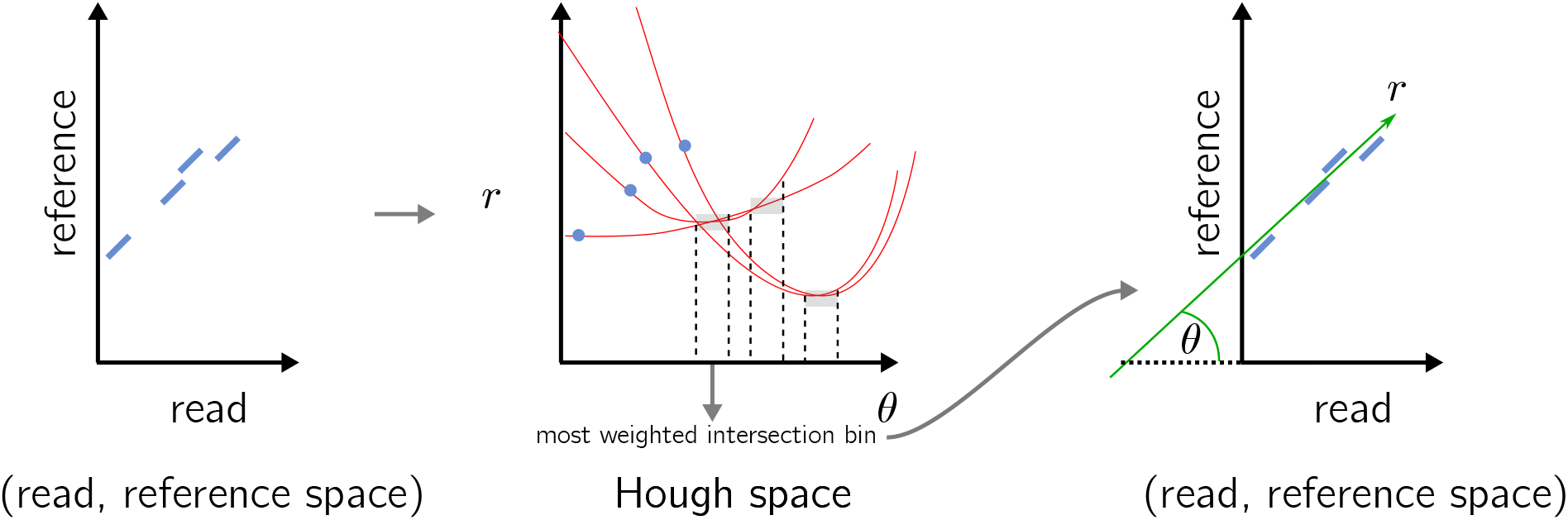
An overview of the Hough transform steps.

## Footnotes

2 and the unpublished `DALIGNER` https://github.com/thegenemyers/DALIGNER

3 https://github.com/bluenote-1577/os-minimap2 and https://github.com/Shamir-Lab/syncmer_mapping

4 https://github.com/waveygang/wfmash/blob/master/README.md, https://github.com/lh3/miniwfa

5 NB: this is different from a de Bruijn graph since nodes with similar contents can be repeated

## References

1 Mohamed Ibrahim Abouelhoda and Enno Ohlebusch. A local chaining algorithm and its applications in comparative genomics. In International Workshop on Algorithms in Bioinformatics, pages 1–16. Springer, 2003.

2 Quim Aguado-Puig, Santiago Marco-Sola, Juan Carlos Moure, Christos Matzoros, David Castells-Rufas, Antonio Espinosa, and Miquel Moreto. Wfa-gpu: Gap-affine pairwise alignment using gpus. bioRxiv, 2022.

3 Mohammed Alser, Jeremy Rotman, Dhrithi Deshpande, Kodi Taraszka, Huwenbo Shi, Pelin Icer Baykal, Harry Taegyun Yang, Victor Xue, Sergey Knyazev, Benjamin D Singer, et al. Technology dictates algorithms: recent developments in read alignment. Genome biology, 22(1):1–34, 2021.

4 Stephen F Altschul, Warren Gish, Webb Miller, Eugene W Myers, and David J Lipman. Basic local alignment search tool. Journal of molecular biology, 215(3):403–410, 1990.

5 Mohammad Ruhul Amin, Steven Skiena, and Michael C Schatz. Nanoblaster: Fast alignment and characterization of oxford nanopore single molecule sequencing reads. In 2016 IEEE 6th International Conference on Computational Advances in Bio and Medical Sciences (ICCABS), pages 1–6. IEEE, 2016.

6 Mahdi Belbasi, Antonio Blanca, Robert S Harris, David Koslicki, and Paul Medvedev. The minimizer jaccard estimator is biased and inconsistent. bioRxiv, 2022.

7 Konstantin Berlin, Sergey Koren, Chen-Shan Chin, James P Drake, Jane M Landolin, and Adam M Phillippy. Assembling large genomes with single-molecule sequencing and locality-sensitive hashing. Nature biotechnology, 33(6):623–630, 2015.

8 Alexander Bowe, Taku Onodera, Kunihiko Sadakane, and Tetsuo Shibuya. Succinct de bruijn graphs. In International workshop on algorithms in bioinformatics, pages 225–235. Springer, 2012.

9 Andrei Z Broder. On the resemblance and containment of documents. In Proceedings. Compression and Complexity of SEQUENCES 1997 (Cat. No. 97TB100171), pages 21–29. IEEE, 1997.

10 Karel Břinda, Maciej Sykulski, and Gregory Kucherov. Spaced seeds improve k-mer-based metagenomic classification. Bioinformatics, 31(22):3584–3592, 07 2015. arXiv:https://academic.oup.com/bioinformatics/article-pdf/31/22/3584/5027960/btv419.pdf, doi: 10.1093/bioinformatics/btv419.

11 Bastien Cazaux, Dmitry Kosolobov, Veli Mäkinen, and Tuukka Norri. Linear time maximum segmentation problems in column stream model. In International Symposium on String Processing and Information Retrieval, pages 322–336. Springer, 2019.

12 Mark J Chaisson and Glenn Tesler. Mapping single molecule sequencing reads using basic localalignment with successive refinement (blasr): application and theory. BMC bioinformatics, 13(1):1–18, 2012.

13 Angana Chakraborty, Burkhard Morgenstern, and Sanghamitra Bandyopadhyay. S-conlsh: Alignment-free gapped mapping of noisy long reads. BMC bioinformatics, 22(1):1–18, 2021.

14 Moses S Charikar. Similarity estimation techniques from rounding algorithms. In Proceedings of the thiry-fourth annual ACM symposium on Theory of computing, pages 380–388, 2002.

15 Chen-Shan Chin and Asif Khalak. Human genome assembly in 100 minutes. BioRxiv, page 705616, 2019.

16 Arthur L Delcher, Simon Kasif, Robert D Fleischmann, Jeremy Peterson, Owen White, andSteven L Salzberg. Alignment of whole genomes. Nucleic acids research, 27(11):2369–2376, 1999.

17 Richard O Duda and Peter E Hart. Use of the hough transformation to detect lines and curves in pictures. Communications of the ACM, 15(1):11–15, 1972.

18 Robert Edgar. Syncmers are more sensitive than minimizers for selecting conserved k-mers in biological sequences. PeerJ, 9:e10805, 2021.

19 Jordan M Eizenga and Benedict Paten. Improving the time and space complexity of the wfa algorithm and generalizing its scoring. bioRxiv, 2022.

20 Marquita Ellis, Giulia Guidi, Aydın Buluç, Leonid Oliker, and Katherine Yelick. dibella: Distributed long read to long read alignment. In Proceedings of the 48th International Conference on Parallel Processing, pages 1–11, 2019.

21 David Eppstein, Zvi Galil, Raffaele Giancarlo, and Giuseppe F Italiano. Sparse dynamic programming ii: convex and concave cost functions. Journal of the ACM (JACM), 39(3):546– 567, 1992.

22 Paolo Ferragina and Giovanni Manzini. Opportunistic data structures with applications. In Proceedings 41st annual symposium on foundations of computer science, pages 390–398. EEE, 2000.

23 Can Firtina, Jisung Park, Jeremie S Kim, Mohammed Alser, Damla Senol Cali, Taha Shahroodi, Nika Mansouri Ghiasi, Gagandeep Singh, Konstantinos Kanellopoulos, Can Alkan, et al. Blend: A fast, memory-efficient, and accurate mechanism to find fuzzy seed matches. arXiv preprint arXiv:2112.08687, 2021.

24 Martin C Frith, Laurent Noé, and Gregory Kucherov. Minimally overlapping words for sequence similarity search. Bioinformatics, 36(22-23):5344–5350, 2020.

25 Yilei Fu, Medhat Mahmoud, Viginesh Vaibhav Muraliraman, Fritz J Sedlazeck, and Todd J Treangen. Vulcan: Improved long-read mapping and structural variant calling via dual-mode alignment. GigaScience, 10(9):giab063, 2021.

26 Zvi Galil and Kunsoo Park. A linear-time algorithm for concave one-dimensional dynamic programming. Information Processing Letters, 1989.

27 Eldar Giladi, John Healy, Gene Myers, Chris Hart, Philipp Kapranov, Doron Lipson, Steve Roels, Edward Thayer, and Stan Letovsky. Error tolerant indexing and alignment of short reads with covering template families. J Comput Biol, 17(10), Oct 2010.

28 Osamu Gotoh. Optimal sequence alignment allowing for long gaps. Bulletin of mathematical biology, 52(3):359–373, 1990.

29 Ehsan Haghshenas, S Cenk Sahinalp, and Faraz Hach. lordfast: sensitive and fast alignment search tool for long noisy read sequencing data. Bioinformatics, 35(1):20–27, 2019.

30 Renmin Han, Yu Li, Xin Gao, and Sheng Wang. An accurate and rapid continuous wavelet dynamic time warping algorithm for end-to-end mapping in ultra-long nanopore sequencing. Bioinformatics, 34(17):i722–i731, 2018.

31 Heikki Hyyrö. A bit-vector algorithm for computing levenshtein and damerau edit distances. Nord. J. Comput., 10(1):29–39, 2003.

32 Lucian Ilie and Silvana Ilie. Multiple spaced seeds for homology search. Bioinformatics, 23(22):2969–2977, 09 2007. arXiv:https://academic.oup.com/bioinformatics/article-pdf/23/22/2969/543804/btm422.pdf, doi:10.1093/bioinformatics/btm422.

33 Silvana Ilie. Efficient computation of spaced seeds. BMC research notes, 5:123–123, 02 2012.

34 Chirag Jain, Alexander Dilthey, Sergey Koren, Srinivas Aluru, and Adam M Phillippy. A fast approximate algorithm for mapping long reads to large reference databases. In International Conference on Research in Computational Molecular Biology, pages 66–81. Springer, 2017.

35 Chirag Jain, Arang Rhie, Nancy F Hansen, Sergey Koren, and Adam M Phillippy. Long-read mapping to repetitive reference sequences using winnowmap2. Nature Methods, pages 1–6, 2022.

36 Chirag Jain, Arang Rhie, Haowen Zhang, Claudia Chu, Brian P Walenz, Sergey Koren, and Adam M Phillippy. Weighted minimizer sampling improves long read mapping. Bioinformatics, 36(Supplement_1):i111–i118, 2020.

37 W James Kent. Blat—the blast-like alignment tool. Genome research, 12(4):656–664, 2002.

38 Szymon M Kiełbasa, Raymond Wan, Kengo Sato, Paul Horton, and Martin C Frith. Adaptive seeds tame genomic sequence comparison. Genome research, 21(3):487–493, 2011.

39 Sam Kovaka, Yunfan Fan, Bohan Ni, Winston Timp, and Michael C Schatz. Targeted nanopore sequencing by real-time mapping of raw electrical signal with uncalled. Nature biotechnology, 39(4):431–441, 2021.

40 Gad M Landau and Uzi Vishkin. Fast parallel and serial approximate string matching. Journal of algorithms, 10(2):157–169, 1989.

41 Roy Lederman. A random-permutations-based approach to fast read alignment. In BMC bioinformatics, volume 14, pages 1–10. BioMed Central, 2013.

42 Heng Li. Aligning sequence reads, clone sequences and assembly contigs with bwa-mem. arXiv preprint arXiv:1303.3997, 2013.

43 Heng Li. Minimap and miniasm: fast mapping and de novo assembly for noisy long sequences. Bioinformatics, 32(14):2103–2110, 2016.

44 Heng Li. Minimap2: pairwise alignment for nucleotide sequences. Bioinformatics, 34(18):3094– 3100, 2018.

45 Heng Li. New strategies to improve minimap2 alignment accuracy. Bioinformatics, 37(23):4572– 4574, 2021.

46 Heng Li, Xiaowen Feng, and Chong Chu. The design and construction of reference pangenome graphs with minigraph. Genome biology, 21(1):1–19, 2020.

47 Ming Li, Bin Ma, Derek Kisman, and John Tromp. Patternhunter ii: highly sensitive and fast homology search. J Bioinform Comput Biol, 2(3):417–439, Sep 2004.

48 Hsin-Nan Lin and Wen-Lian Hsu. Kart: a divide-and-conquer algorithm for ngs read alignment. Bioinformatics, 33(15):2281–2287, 2017.

49 Bo Liu, Yan Gao, and Yadong Wang. Lamsa: fast split read alignment with long approximate matches. Bioinformatics, 33(2):192–201, 2017.

50 Bo Liu, Dengfeng Guan, Mingxiang Teng, and Yadong Wang. rHAT: fast alignment of noisy long reads with regional hashing. Bioinformatics, 32(11):1625–1631, 11 2015. arXiv:https://academic.oup.com/bioinformatics/article-pdf/32/11/1625/22645531/btv662.pdf, doi: 10.1093/bioinformatics/btv662.

51 Bo Liu, Yadong Liu, Junyi Li, Hongzhe Guo, Tianyi Zang, and Yadong Wang. desalt: fast and accurate long transcriptomic read alignment with de bruijn graph-based index. Genome biology, 20(1):1–14, 2019.

52 Santiago Marco-Sola, Jordan M Eizenga, Andrea Guarracino, Benedict Paten, Erik Garrison, and Miquel Moreto. Optimal gap-affine alignment in o (s) space. bioRxiv, 2022.

53 Santiago Marco-Sola, Juan Carlos Moure, Miquel Moreto, and Antonio Espinosa. Fast gap-affine pairwise alignment using the wavefront algorithm. Bioinformatics, 37(4):456–463, 2021.

54 Josip Marić, Ivan Sović, Krešimir Križanović, Niranjan Nagarajan, and Mile Šikić. Graphmap2-splice-aware rna-seq mapper for long reads. bioRxiv, page 720458, 2019.

55 Frédéric Meunier, Olivier Gandouet, Éric Fusy, and Philippe Flajolet. Hyperloglog: the analysis of a near-optimal cardinality estimation algorithm. Discrete Mathematics & Theoretical Computer Science, 2007.

56 Alla Mikheenko, Andrey V Bzikadze, Alexey Gurevich, Karen H Miga, and Pavel A Pevzner. Tandemtools: mapping long reads and assessing/improving assembly quality in extra-long tandem repeats. Bioinformatics, 36(Supplement_1):i75–i83, 2020.

57 Hamid Mohamadi, Justin Chu, Benjamin P Vandervalk, and Inanc Birol. nthash: recursive nucleotide hashing. Bioinformatics, 32(22):3492–3494, 2016.

58 Gene Myers. A fast bit-vector algorithm for approximate string matching based on dynamic programming. Journal of the ACM (JACM), 46(3):395–415, 1999.

59 Saul B Needleman and Christian D Wunsch. A general method applicable to the search for similarities in the amino acid sequence of two proteins. Journal of molecular biology, 48(3):443–453, 1970.

60 Christian Otto, Steve Hoffmann, Jan Gorodkin, and Peter F Stadler. Fast local fragment chaining using sum-of-pair gap costs. Algorithms for Molecular Biology, 6(1):1–8, 2011.

61 David Pellow, Abhinav Dutta, and Ron Shamir. Using syncmers improves long-read mapping. bioRxiv, 2022.

62 Jingwen Ren and Mark JP Chaisson. lra: A long read aligner for sequences and contigs. PLOSComputational Biology, 17(6):e1009078, 2021.

63 Michael Roberts, Wayne Hayes, Brian R Hunt, Stephen M Mount, and James A Yorke. Reducing storage requirements for biological sequence comparison. Bioinformatics, 20(18):3363– 3369, 2004.

64 Kristoffer Sahlin. Effective sequence similarity detection with strobemers. Genome research, 31(11):2080–2094, 2021.

65 Kristoffer Sahlin. Faster short-read mapping with strobemer seeds in syncmer space. bioRxiv, 2021.

66 Kristoffer Sahlin and Veli Mäkinen. Accurate spliced alignment of long rna sequencing reads. Bioinformatics, 37(24):4643–4651, 2021.

67 Kristoffer Sahlin and Paul Medvedev. Error correction enables use of oxford nanopore technology for reference-free transcriptome analysis. Nature communications, 12(1):1–13, 2021.

68 Saul Schleimer, Daniel S Wilkerson, and Alex Aiken. Winnowing: local algorithms for document fingerprinting. In Proceedings of the 2003 ACM SIGMOD international conference on Management of data, pages 76–85, 2003.

69 Fritz J Sedlazeck, Philipp Rescheneder, Moritz Smolka, Han Fang, Maria Nattestad, Arndt Von Haeseler, and Michael C Schatz. Accurate detection of complex structural variations using single-molecule sequencing. Nature methods, 15(6):461–468, 2018.

70 Kishwar Shafin, Trevor Pesout, Ryan Lorig-Roach, Marina Haukness, Hugh E Olsen, Colleen Bosworth, Joel Armstrong, Kristof Tigyi, Nicholas Maurer, Sergey Koren, et al. Nanopore sequencing and the shasta toolkit enable efficient de novo assembly of eleven human genomes. Nature biotechnology, 38(9):1044–1053, 2020.

71 Jim Shaw and Yun William Yu. Theory of local k-mer selection with applications to long-read alignment. bioRxiv, 2021.

72 Ivan Sović, Mile Šikić, Andreas Wilm, Shannon Nicole Fenlon, Swaine Chen, and Niranjan Nagarajan. Fast and sensitive mapping of nanopore sequencing reads with graphmap. Nature communications, 7(1):1–11, 2016.

73 Esko Ukkonen. Algorithms for approximate string matching. Information and control, 64(1-3):100–118, 1985.

74 Ze-Gang Wei, Xing-Guo Fan, Hao Zhang, Xiao-Dan Zhang, Fei Liu, Yu Qian, and Shao-Wu Zhang. kngmap: sensitive and fast mapping algorithm for noisy long reads based on the k-mer neighborhood graph. Frontiers in Genetics, page 988, 2022.

75 Thomas D Wu and Colin K Watanabe. Gmap: a genomic mapping and alignment program for mrna and est sequences. Bioinformatics, 21(9):1859–1875, 2005.

76 Chuan-Le Xiao, Ying Chen, Shang-Qian Xie, Kai-Ning Chen, Yan Wang, Yue Han, Feng Luo, and Zhi Xie. Mecat: fast mapping, error correction, and de novo assembly for single-molecule sequencing reads. nature methods, 14(11):1072–1074, 2017.

77 Haowen Zhang, Haoran Li, Chirag Jain, Haoyu Cheng, Kin Fai Au, Heng Li, and Srinivas Aluru. Real-time mapping of nanopore raw signals. Bioinformatics, 37(Supplement_1):i477–i483, 2021.

